# CREB5 reprograms nuclear interactions to promote resistance to androgen receptor targeting therapies

**DOI:** 10.1101/2021.08.18.456892

**Authors:** Justin Hwang, Rand Arafeh, Ji-Heui Seo, Sylvan C. Baca, Megan Ludwig, Taylor E. Arnoff, Camden Richter, Hannah E. Bergom, Sean McSweeney, Jonathan P. Rennhack, Sarah A. Klingenberg, Alexander TM. Cheung, Jason Kwon, Jonathan So, Steven Kregel, Eliezer M. Van Allen, Justin M. Drake, Mathew L. Freedman, William C. Hahn

## Abstract

Metastatic castration resistant prostate cancers (mCRPC) are treated with therapies that antagonize the androgen receptor (AR). Nearly all patients develop resistance to AR-targeted therapies (ART). Our previous work identified CREB5 as an upregulated target gene in human mCRPC that promoted resistance to all clinically-approved ART. The mechanisms by which CREB5 promotes progression of mCRPC or other cancers remains elusive. Integrating ChIP-seq and rapid immunoprecipitation and mass spectroscopy of endogenous proteins (RIME), we report that cells overexpressing CREB5 demonstrate extensive reprogramming of nuclear protein-protein interactions in response to the ART agent enzalutamide. Specifically, CREB5 physically interacts with AR, the pioneering actor FOXA1, and other known co-factors of AR and FOXA1 at transcription regulatory elements recently found to be active in mCRPC patients. We identified a subset of CREB5/FOXA1 co-interacting nuclear factors that have critical functions for AR transcription (GRHL2, HOXB13) while others (TBX3, NFIC) regulated cell viability and ART resistance and were amplified or overexpressed in mCRPC. Upon examining the nuclear protein interactions and the impact of CREB5 expression on the mCRPC patient transcriptome, we found CREB5 was associated with TGFβ and Wnt signaling and epithelial to mesenchymal transitions, implicating these pathways in ART resistance. Overall, these observations define the molecular interactions among CREB5, FOXA1, and pathways that promote ART resistance.

## Introduction

The androgen receptor (AR) plays a fundamental role in the development and function of the prostate. AR transcriptional activity is required for the initiation and progression of prostate cancer and remains critical in metastatic castration resistant prostate cancer (mCRPC) (Abida et al., 2019; Armenia et al., 2018; Grasso et al., 2012; He et al., 2021; Robinson et al., 2015). The standard of care for patients with mCRPC is androgen deprivation therapy (ADT) and AR-targeted therapies (ART). While ADT and ART initially induce responses and lengthen the survival of mCRPC patients, intrinsic or acquired resistance continues to be a substantial clinical obstacle. With limited treatment options, better mechanistic understanding of the targets that drive resistant disease are urgently needed.

Several lines of evidence indicate that mCRPC may acquire resistance to ART through multiple mechanisms. Recent studies of mCRPC patients resistant to second generation ART and ADT have identified that a subset of the resistant mCRPC harbor AR genomic alterations through mutations, copy number gain, enhancer amplification, as well as increased resistant transcript variants (ARV7) (Bubley and Balk, 2017; Henzler et al., 2016; Takeda et al., 2018; Viswanathan et al., 2018). Other clinical and functional studies together demonstrate that resistance also occurs through mechanisms including upregulation of cell cycle genes (Comstock et al., 2013; Han et al., 2017), and/or the proto-oncogene MYC (Abida et al., 2019; Armenia et al., 2018; Bernard et al., 2003; Grasso et al., 2012; Robinson et al., 2015; Sharma et al., 2013). In addition, selective activation of pathways such as epithelial to mesenchymal transition, TGFβ and Wnt have been recently identified from profiling mCRPC patient become resistant to second generation ART or ADT (Alumkal et al., 2020; He et al., 2021).

Recent mechanistic studies have begun to elucidate how dysregulated epigenomic and transcriptomic processes drive disease pathogenesis and progression. AR activity in prostate tissue is regulated by AR co-factors such as HOXB13 and FOXA1 at AR binding sites (ARBs) (Pomerantz et al., 2015; Pomerantz et al., 2020). ChIP-seq experiments have demonstrated that FOXA1, a co-factor with both pioneering and transcription functions, binds to distinct sites and has different protein interactions that is dependent on the disease stage and pathological phenotype (Baca et al., 2021; Pomerantz et al., 2015; Pomerantz et al., 2020). Recent studies indicate that genomic alterations of *FOXA1* promote ART resistance (Adams et al., 2019; Baca et al., 2021; Parolia et al., 2019; Shah and Brown, 2019). We have shown FOXA1 function remains functionally relevant even as mCRPC differentiate into aggressive variants that no longer require AR signaling, such as neuroendocrine prostate cancer (Baca et al., 2021). We have previously demonstrated that the transcriptional factor cAMP responsive element binding protein 5 (C*REB5*) promoted resistance to FDA-approved ART, enzalutamide, darolutamide and apalutamide (Hwang et al., 2019). CREB5 has also been associated with progression of cancers including ovarian, colorectal and breast (Bhardwaj et al., 2017; He et al., 2017; Molnar et al., 2018; Qi and Ding, 2014). However, the molecular mechanisms by which CREB5 promotes resistance in mCRPC or general tumorigenesis remains unclear.

Here we utilized ChIP-seq and rapid immunoprecipitation and mass spectrometry of endogenous proteins (RIME) experiments to resolve the CREB5 associated transcriptional targets and molecular interactions. These CREB5 transcriptional co-factors are potential therapeutic targets to perturb CREB5 signaling in cancers that upregulate its activity.

## Materials and Methods

### Genome-scale ORF screen analysis

We analyzed a published genome-scale ORF screen performed in LNCaP cells (Hwang et al., 2019). Specifically, we compared the experimental arms conducted in control media (FCS) and low androgen media (CSS) containing enzalutamide. Z-scores represent the relative effects of each ORF on cell proliferation after 25 days in culture.

### Rapid Immunoprecipitation Mass Spectrometry of Endogenous Proteins (RIME)

Upon preparation of cells in each experimental arm, two replicates of 50 million cells each were fixed in 1% formaldehyde for 10 minutes. Reactions were terminated in 0.125M glycine for 5 minutes. Cells were subsequently collected and lysed at 4°C in RIPA buffer (Cell Signaling Technology, 13202S) containing protease inhibitor cocktail (Roche, 11836145001). Cells were then sonicated with a Covaris sonicator to yield DNA fragments averaging around 3000 nucleotides. To target V5 tagged CREB5, CREB5 L434P, or CREB5, 20μL of V5 antibody (Cell Signaling Technology, 58613) was added to the supernatant. To target FOXA1, each sample was incubated with 20μL of two FOXA1 antibodies against distinct epitopes (Abcam, ab23738 and Cell Signaling Technology, 58613). Samples were incubated with gentle mixing at 4°C overnight. The following morning, RIPA buffer was used to wash Protein A magnetic beads (Life Technologies, 88846) 5 times, and the beads were then resuspended into the original volume. 50μL of the bead mixture was added to each sample and incubated for 2 hours at 4°C. Each sample was then washed 5 times with 300μL of RIPA buffer, followed by 5 times with 300μL of Ammonium Bicarbonate (50μM), and finally resuspended in 50μL of Ammonium Bicarbonate (50μM). These samples on the beads were then sent for proteomic analysis at the Taplin Mass Spectrometry Facility at Harvard Medical School. Upon obtaining mapped reads, only unique peptides of proteins were considered for subsequent analysis. Beads were subjected to trypsin digestion procedure (Shevchenko et al., 1996) then washed and dehydrated with acetonitrile for 10 min followed by removal of acetonitrile. The beads were then completely dried in a speed-vac. Rehydration of the beads was with 50 mM ammonium bicarbonate solution containing 12.5 ng/µl modified sequencing-grade trypsin (Promega, Madison, WI) at 4°C. After 45 min., the excess trypsin solution was removed and replaced with 50 mM ammonium bicarbonate solution to just cover the gel pieces. Samples were then placed in a 37°C room overnight. Peptides were later extracted by removing the ammonium bicarbonate solution, followed by one wash with a solution containing 50% acetonitrile and 1% formic acid. The extracts were then dried in a speed-vac (∼1 hr). The samples were then stored at 4°C until analysis. On the day of analysis the samples were reconstituted in 5 - 10 µl of HPLC solvent A (2.5% acetonitrile, 0.1% formic acid). A nano-scale reverse-phase HPLC capillary column was created by packing 2.6 µm C18 spherical silica beads into a fused silica capillary (100 µm inner diameter x ∼30 cm length) with a flame-drawn tip (Peng and Gygi, 2001). After equilibrating the column each sample was loaded via a Famos auto sampler (LC Packings, San Francisco CA) onto the column. A gradient was formed and peptides were eluted with increasing concentrations of solvent B (97.5% acetonitrile, 0.1% formic acid). As peptides eluted they were subjected to electrospray ionization and then entered into an LTQ Orbitrap Velos Pro ion-trap mass spectrometer (Thermo Fisher Scientific, Waltham, MA). Peptides were detected, isolated, and fragmented to produce a tandem mass spectrum of specific fragment ions for each peptide. Peptide sequences (and hence protein identity) were determined by matching protein databases with the acquired fragmentation pattern by the software program, Sequest (Thermo Fisher Scientific, Waltham, MA) (Eng et al., 1994). All databases include a reversed version of all the sequences and the data was filtered to between a one and two percent peptide false discovery rate.

### Population Doubling

100∼200k cells were plated in 12-well plates in either control media (FCS) or low androgen media (CSS) with 10μM of enzalutamide. Cell counts and relative cell viability were determined after 7 days using a Vi-Cell. Original cell counts were subtracted before doubling was computed.

### Generation of CREB5 Point Mutants

A pDNR221 CREB5 plasmid was used as a backbone for point mutagenesis. Per the five point mutants, forward and reverse primers were designed to mutagenize nucleotides of CREB5 to reflect the corresponding changes in protein sequence. Primers were designed with primerX (http://www.bioinformatics.org/primerx). Detailed reaction conditions were followed according to a Quikchange II Site-Directed Mutagenesis Kit (Agilent, 2005230). The mutant clones were each sequenced to confirm their identities before subsequent use in recombination reactions. LR Clonase II (Invitrogen, 11791-020) was used to catalyze the recombination reactions to insert each mutated CREB5 ORF into a puromycin resistant lentiviral pLX307 vector. Positive clones were then sequenced to confirm the identity of the resulting mutant vectors used for further experimentation.

### CRISPR-Cas9 experiments

To generates sgRNAs, oligos were cloned into a pXPR_003 vector as previously cited (Hwang et al., 2019). Blasticidin-resistant Cas9-positive LNCaP enzalutamide resistant cells were cultured in 10 μg/ml blasticidin (Thermo Fisher Scientific, NC9016621) for 72 h to select for cells with optimal Cas9 activity. One million cells were seeded in parallel in 6-well plates and infected with lentiviruses expressing puromycin-resistant sgRNAs targeting FOXA1, TBX3, NFIC or GFP control. After 48h, cells were counted and seeded, using a Vi-Cell, with 20 uM enzalutamide at a density of 20 000 cells per well in 6 well plate for a proliferation assay. After 24h, cells were subjected to puromycin selection for 3 days. 7 days later, cells were counted again with a Vi-Cell to assess viability, representing a total of 12 days. The target sequences against GFP were AGCTGGACGGCGACGTAAA (sgGFP1) and GCCACAAGTTCAGC GTGTCG (sgGFP2). The target sequences against FOXA1 were GTTGGACGGC GCGTACGCCA (sgFOXA1-1), GTAGTAGCTGTTCCAGTCGC (sgFOXA1-2), and ACTGCGCCCCCCATA AGCTC (sgFOXA1-4). The target sequences against TBX3 were GAAAAGGTGAGCCTTGACCG (sgTBX3-1), and GCTCTTACAATGTGGAACCG (sgTBX3-2). The target sequences against NFIC were ACGGCCACGCCAATGTGGTG (sgNFIC-1), and GCTGAGCATCACCGGCAAGA (sgNFIC-2).

### Immunoblotting

Cells were lysed using 2 × sample buffer (62.5 mM Tris pH 6.8, 2% SDS, 10% Glycerol, Coomassie dye) and freshly added 4% β-mercaptoethanol. Lysed cells were scraped, transferred into a 1.5 mL microcentrifuge tube, sonicated for 15 seconds and boiled at 95C for 10 minutes. Proteins were resolved in NuPAGE 4-12% Bis-Tris Protein gels (Thermo Fisher Scientific) and run with NuPAGE MOPS SDS Running Buffer (Thermo Fisher Scientific, NP0001). Proteins were transferred to nitrocellulose membranes using an iBlot apparatus (Thermo Fisher Scientific). Membranes were blocked in Odyssey Blocking Buffer (LI-COR Biosciences, 927-70010) for one hour at room temperature, and membranes were then cut and incubated in primary antibodies diluted in Odyssey Blocking Buffer at 4°C overnight. The following morning, membranes were washed with Phosphate-Buffer Saline, 0.1% Tween (PBST) and incubated with fluorescent anti-rabbit or anti-mouse secondary antibodies at a dilution of 1:5000 (Thermo Fisher Scientific, NC9401842 (Rabbit) and NC0046410 (mouse)) for one hour at room temperature. Membranes were again washed with PBST and then imaged using an Odyssey Imaging System (LI-COR Biosciences). Primary antibodies used include V5 (Cell Signaling Technology, 13202S), β-actin (Cell Signaling Technology, 8457L), TBX3 (Life Technologies, 424800), NFIC (Sigma, SAB2101580), Tubulin (Sigma, T9026).

### Overlap analysis of CREB5 binding sites

Bed files containing peak summit locations from determined by MACS2 from CREB5 ChIP-seq data were intersected with the indicates datasets using BEDtools v2.27.1 (Quinlan and Hall, 2010). To assess overlap with FOXA1 binding sites, FOXA1 ChIP-seq datasets from 23 prostate adenocarcinoma patient-derived xenografts (Nguyen et al., 2017) and tissue (Pomerantz et al., 2015; Pomerantz et al., 2020) were merged to create a FOXA1 union peak set. To assess overlap of CREB5 binding and various sets of AR binding sites, we first created a union set of CREB5 peaks that were present with or without enzalutamide treatment. This CREB5 peak sets was intersected with the indicated AR peaks sets (**Figure 2B**) from references (Pomerantz et al., 2015; Pomerantz et al., 2020). The percentage of each class of AR+CREB5+ peaks were assessed for overlap with the union set of FOXA1 peaks in Fig 2C.

### Project Achilles 2.20.1 analysis

Of the total of 503 cell lines, we analyzed a published genome-scale RNAi screen of 8 prostate cancer cell lines (Cowley et al., 2014; Shalem et al., 2014; Tsherniak et al., 2017) whereby we averaged the dependency for each gene. Cell lines included NCIH660 (NEPC-like), PC3 and DU145 (AR negative), 22RV1 (expressing an AR V7 splice variant), LNCaP, VCaP, and MDAPCA2B (AR positive and dependent), and PRECLH (normal immortalized prostate epithelium).

### Motif enrichment analysis

Known motifs enriched in TBX3 and NFIC ChIP-seq data from HepG2 cells (GEO: GSM2825557, GSM2902642) compared to a whole-genome background were identified with Homer version 4.17 (Heinz et al., 2010). Selected examples from the most significantly enriched known motifs are shown (**Figure 4D**).

### Expression association analysis of CREB5 in mCRPC

We analyzed RNA-sequencing data from an updated combined cohort of men with mCRPC from multiple institutions comprising the SU2C/PCF Prostate Cancer Dream Team (Abida et al., 2019). RNA-seq data, normalized in units of transcripts per million (TPM), was available from 208 patients. Expression data was previously examined and adjusted for batch effects using ComBat (Johnson et al., 2007) via the R Bioconductor package “sva” (Leek et al., 2012), version V3.22.0. The Spearman correlations were determined for CREB5 against all detectable transcripts in these samples. This profile of association was further examined with focus on CREB5 and its associate with genes in the indicated pathways using pre-defined MSigDB signatures.

## Results

### CREB5 drives a resistance response to enzalutamide and androgen deprivation

We previously performed a large-scale screen to identify genes involved in ADT and ART resistance through overexpression of 17,255 ORFs in LNCaP cells, an AR-dependent prostate cancer cell line (Hwang et al., 2019). We and others functionally validated several genes that drive ADT/ART resistance (FGFs, CDK4/6, MDM4, CREB5) in cell lines or mCRPC patients (Bluemn et al., 2017; Comstock et al., 2013; Elmarakeby et al., 2020; Han et al., 2017; Hwang et al., 2019). Here we sought to interrogate the molecular mechanisms specifically associated with resistance to ADT and the clinically used ART, enzalutamide. Specifically, we identified ORFs that when overexpressed only promoted cell survival in the presence of ADT/ART and not in standard cultures. Upon re-examining the data, we found that unlike in ADT/ART, overexpression of CREB5 reduced viability (Z= -1.3) in standard cultures (**Figure 1A**). The Z-scores, which represent relative proliferation effects compared to all 17,255 ORFs exhibited robust differentials when comparing the treated arm in comparison to the control arm (Z = +14.5 and -1.3). This observation contrasted to other genes that mediate resistance to ART, such as CDK4 or CDK6, which promoted cell fitness regardless of treatment conditions. At genome scale, many ORFs had preferential fitness effects when considering the differential Z-score of the treated and standard conditions. Among ORFs, CREB5 had the greatest differential viability effect after low androgen and enzalutamide treatment (**Supplemental Table 1**). Overall, these observations prompted us to pursue a deeper interrogation of binding properties of CREB5 to understand specific molecular interactions that promote resistance to ART.

**Figure 1.**
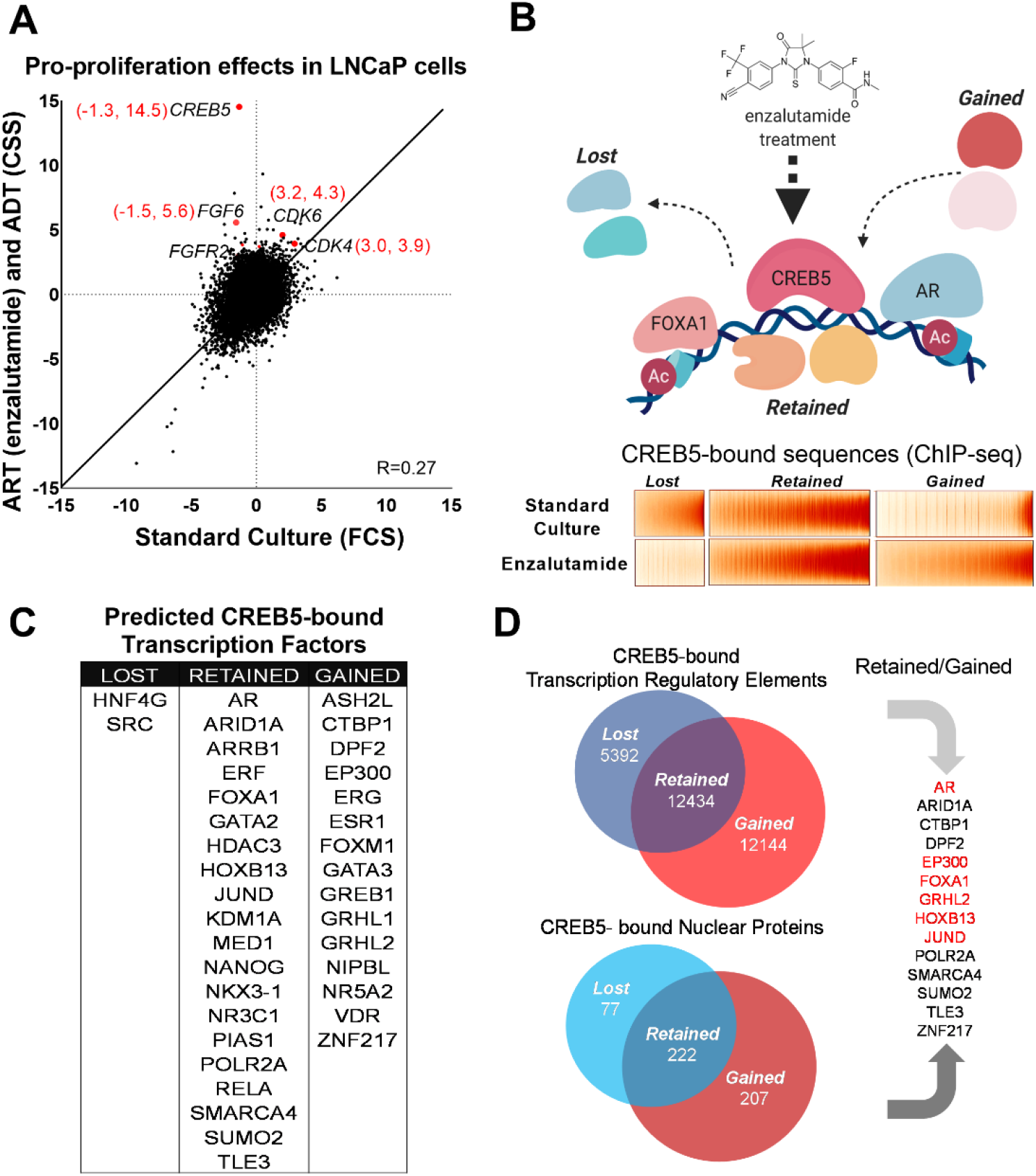
CREB5 overexpression and nuclear interactions that are reprogrammed upon ART treatments. (A) Analysis of enzalutamide resistance genes in LNCaP cells based on a genome-scale screen including 17,255 ORFs. Z-scores are displayed for the experimental arms conducted in either standard culture (FCS, x-axis) or treatment (enzalutamide + CSS, y-axis) conditions. *CREB5* and other enzalutamide specific hits (Z>3) and their proliferation scores are highlighted in red. (B) A model that depicts changes in chromatin associated interactions of CREB5 that occur post enzalutamide treatment. Bottom, CREB5 ChIP-seq data is presented in accordance to three categories of CREB5 binding behavior. Categories are grouped by significant changes by enzalutamide treatments. (C) GIGGLE analyses predicts transcription factors that are CREB5-bound based on the ChIP-seq experiments as categorized in B). (D) RIME experiments were performed to identify CREB5 interaction profiles in control or enzalutamide treated cultures. The common proteins identified by both RIME and GIGGLE are highlighted for the retained and gained groups.

### Dynamic CREB5 nuclear interactions are associated with the ART resistance response

We next determined the CREB5 cistrome as well as other regulatory proteins critical to the ART resistant phenotype by identifying features that were either “retained” or “gained” upon ART treatments. In LNCaP cells overexpressing V5-tagged CREB5 or luciferase and cultured in either standard media or media containing enzalutamide, we analyzed differential interactions of CREB5 at regulatory elements through ChIP-seq or with other proteins through RIME. We first examined CREB5 binding sites pre- and post-enzalutamide treatments by performing CREB5 ChIP-seq. CREB5 overexpression induced a robust differential phenotype (**Figure 1B**), in which lost (n = 5392), retained (n = 12,432), or gained (n = 12,144) CREB5 binding sites were tallied in the enzalutamide condition compared to the pre-treatment condition. To nominate other candidate trans-acting factors that bind within the three defined categories, we used the GIGGLE, an analytical approach that compares collection of sequences from ChIP-seq experiments with over 10,000 experiments from the ENCODE ChIP-seq database and nominates the transcription factors with highly similar binding profiles (Layer et al., 2018). We observed that the elements in which CREB5 retained or gained sites were statistically significantly enriched of other nuclear proteins in which ChIP-seq had been performed in prostate cancer cell lines. This included binding elements (AR, FOXA1, HOXB13) that we previously demonstrated through ChIP-seq experiments (**Figure 1C**). In addition, we also identify that retained or gained sequences nominated well studied regulators or co-factors of AR such as GRHL2 (Paltoglou et al., 2017), EP300 (Yu et al., 2020), and SMARCA4 (Launonen et al., 2021; Marshall et al., 2003). These observations suggested that CREB5 bound critical regulators of prostate cancer biology to drive ART resistance.

Rapid immunoprecipitation mass spectrometry of endogenous proteins (RIME) has been used as a tool to study transcription co-factors interactions in hormone regulated cancer cells (Glont et al., 2019; Mohammed et al., 2016). To build upon the predictions of CREB5 binding by GIGGLE, we directly examined CREB5 nuclear interactions via RIME. As part of the RIME analysis, we included only the unique peptides that bound CREB5 after subtraction of all proteins precipitated by the V5 antibody or luciferase in cell lysates (**Supplemental Table 2**). When we compared the analyzed RIME binding profiles of the control to the enzalutamide treated conditions, analogous to the ChIP-seq experiments, we found the CREB5-bound unique peptides also segregated into lost (n=77), retained (n=222) and gained groups (n=207) (**Figure 1D**). To consider key factors in the retained or gained groups that associate with resistance, we integrated the results derived from GIGGLE and RIME. From this we found both approaches nominated known AR interactions (EP300, FOXA1, and GRHL2). Overall, these parallel approaches demonstrate that CREB5 binding dynamically responds to ART treatment. Moreover, the retained or gained RIME interactions indicate CREB5 may interact with distinct sets of co-factors, some of which are AR associated, to promote ART resistance.

### CREB5-FOXA1 interactions converge in ART resistance

In LNCaP cells, we comprehensively compared CREB5 and FOXA1 protein interactions using RIME. In prior work in LNCaP cells, we found that overexpressing CREB3, a related CREB family member, conferred significantly weaker resistance to enzalutamide, and therefore serves as a useful control to understand CREB5 functions in resistance (Hwang et al., 2019). We used RIME to target overexpressed V5-tagged CREB5, luciferase, or CREB3 upon treating cells with enzalutamide. We optimized experimental conditions to consistently observe unique peptides representing CREB5, CREB3, and FOXA1. On average, we detected 8, 23, and 14 unique peptides that respectively mapped to CREB5, CREB3, and FOXA1 (**Supplemental Table 3**). RIME interaction profiles were subsequently constructed for each targeted protein based on the visualization of the counts of all detected unique peptides that were bound. The overall RIME profiles of CREB5 and FOXA1 were compared and exhibited a positive correlation (R= 0.394), while those of CREB3 with either CREB5 or FOXA1 lacked correlation (R= 0.0332, -0.0498) (**Figure 2A**). At the peptide level, we found that CREB5 and FOXA1 shared a total of 504 protein interactions at chromatin and of these 504, 335 did not interact with CREB3. While almost three times the number of CREB3 peptides were detected relative to CREB5, CREB3 and FOXA1 shared only 83 unique interactions (**Figure 2B**). These observations nominated CREB5/FOXA1 specific protein-protein interactions.

**Figure 2.**
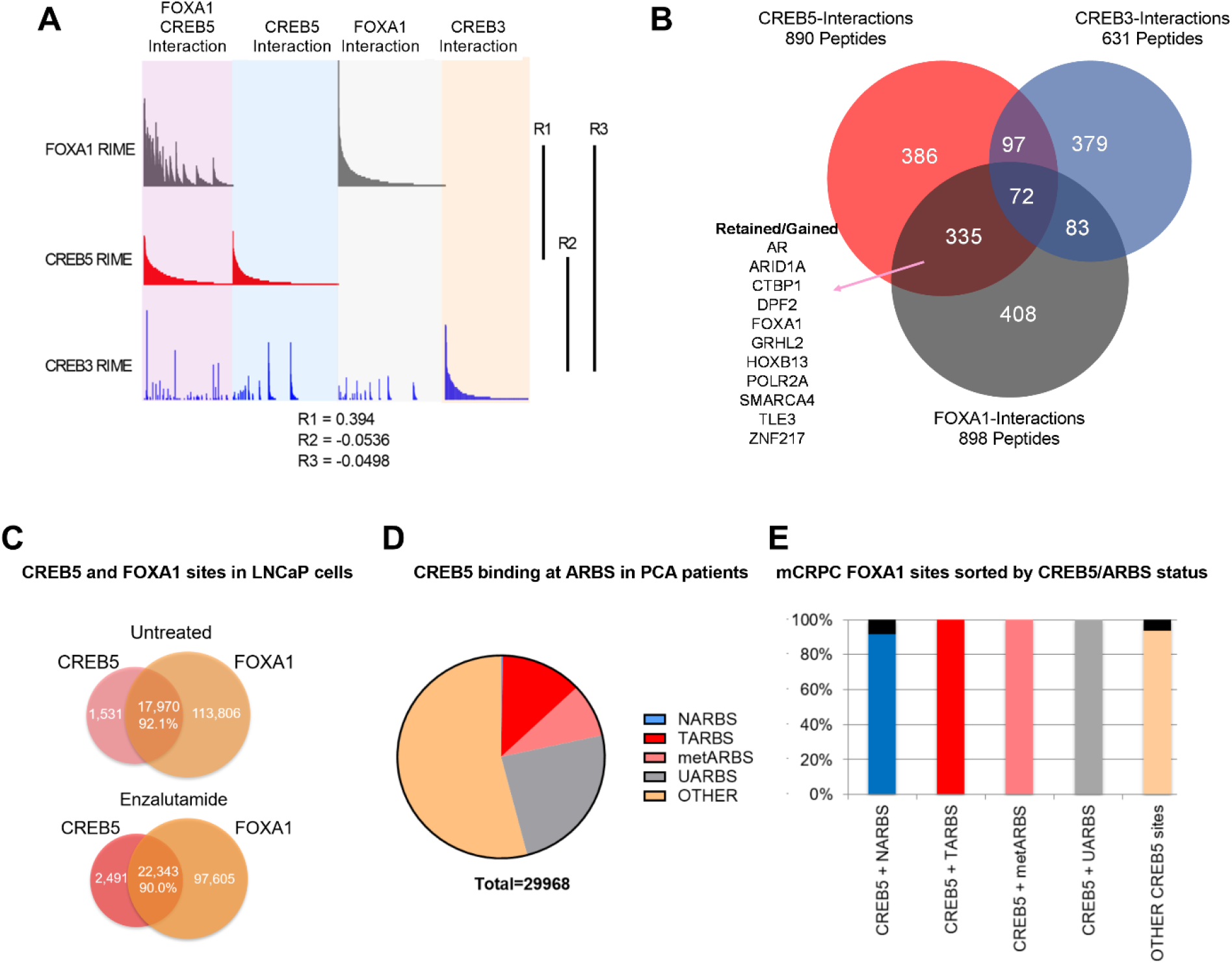
CREB5 and FOXA1 share chromatin associated functions in mCRPC based on binding sequences and RIME interaction profiles. (A) RIME analysis depicting the interaction profiles of FOXA1 (grey), CREB5 (red), CREB3 (blue). Proteins that interact with FOXA1 and CREB5 are also shown. The Pearson correlation coefficients (R) are shown (B) Venn diagram depicting unique peptide interactions that are either independent or shared between CREB5 (red), CREB3 (blue), and FOXA1 (grey). Peptides identified to be induced by enzalutamide are highlighted as Retained/Gained. (C) ChIP-seq experiments were used to examine CREB5 and FOXA1 interactions in LNCaP cells with or without enzalutamide treatments. The VENN diagram depict total binding sites in each condition and the overlapping sites and percentage of shared transcription regulatory elements. (D) CREB5 bound sites are analyzed and represented as AR binding sites (ARBS) observed in clinical samples. This includes ARBS exclusive in normal (NARBS), tumor (TARBS), mCRPC (metARBS), all tissues (UARBS), as well as all non ARBS (OTHER). (E) CREB5 bound ARBs are further classified and depicted as % of FOXA1 sites observed in mCRPC (y-axis). The colors represent the overall percentage of FOXA1 sites while the black represents non FOXA1 sites.

As an orthogonal approach to RIME we sought to examine ChIP-seq interactions of CREB5-FOXA1. We first examined the overlap of CREB5 and FOXA1 at transcription regulatory elements with and without ART. By analyzing ChIP-seq experiments that targeted either CREB5 or FOXA1, we found that regardless of ART, CREB5 and FOXA1 shared a strong degree of interactions as more than 90% of CREB5 bound sites were FOXA1 bound (**Figure 2C**). Pomerantz et al. have recently demonstrated that AR binding sites are reprogrammed during tumorigenesis and progression (Pomerantz et al., 2015; Pomerantz et al., 2020). Since CREB5 promoted ART resistant activity, we anticipated CREB5 interactions at ARBs would be enriched in tumor specific binding sites. When considering the subset of ARBs specific to progression, CREB5 indeed bound ARBs in prostate cancer (12.7%) or mCRPC (8.6%) tissue at higher rates as compared to normal prostate ARBs (0.2%) (**Figure 2D**). Unlike ARBs, FOXA1 binding was less dynamic and the binding sites were relatively ubiquitous in prostate tissue. Almost all CREB5 bound sites were also observed in mCRPCs and were FOXA1 bound (**Figure 2E**). Taken together these observations demonstrate that CREB5 and FOXA1 engage a similar subset of proteins in cells with ART resistance and these interactions are enriched at ARBs or FOXA1 binding sites in mCRPC.

### CREB5 interacting co-factors are associated with ART resistance

To define which CREB5-specific interactions are required to drive ART resistance, we used CREB5 mutants to interrogate these interactions. Based on sequence alignment, the B-Zip and L-Zip domains are highly homologous in CREB and ATF family members (**Figure 3A**) and regulate binding to DNA- and CREB co-factors (Dwarki et al., 1990; Luo et al., 2012). Within the B-Zip and L-Zip domains, several leucine residues regulate transcriptional activity and heterodimerization with the transcription factors JUN and FOS *in vitro* (Fuchs and Ronai, 1999; Nomura et al., 1993). We engineered CREB5 point mutants that would emulate these structural perturbations by disrupting binding at chromatin (R396E), CRTCs (K405A and K406A), and JUN/FOS (L431P and L434P). We expressed these CREB5 variants in LNCaP cells (**Figure 3B**) and found that despite robust expression of L431P and L434P CREB5, cells expressing these mutants proliferated similar to cells expressing luciferase controls in cell doubling assays performed in low androgen media containing enzalutamide (**Figure 3B**), indicating L431P and L434P CREB5 lacked interactions critical for ART resistance.

**Figure 3.**
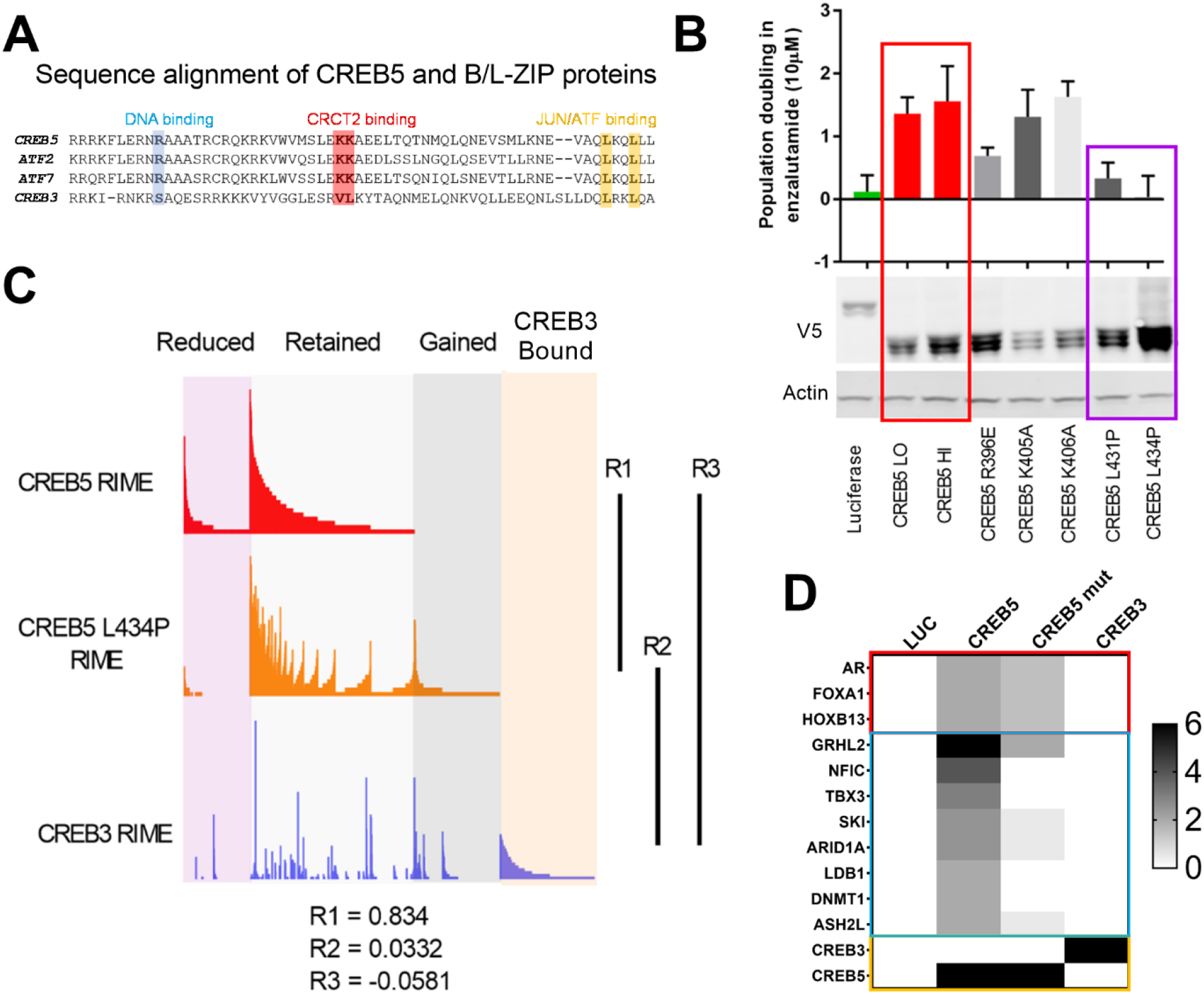
A loss of resistant CREB5 mutant was identified and determines transcription co-regulators associated with ART resistant proliferation. (A) Alignment of CREB5 sequence with ATF2, ATF7, and CREB3, highlighting the DNA binding domains (blue), CRCT2 binding domains (red), and JUN/ATF binding domains (yellow). (B) Population doubling (y-axis) of LNCaP cells overexpressing wild-type CREB5 variants (red), CREB5 JUN/FOS-binding mutants (purple), and a luciferase negative control (green) in 10μM enzalutamide. V5 expression represents V5-tagged CREB5 protein levels. Actin is a loading control. (C) RIME analysis depicting the interaction profiles of wild-type CREB5 (red), CREB5 L434P (orange), and CREB3 (blue). CREB5 interactions that were reduced, retained, or gained upon enzalutamide treatments are depicted. The Pearson correlation coefficients (R) are shown. (D) A heatmap depicts the RIME interactions of luciferase control, wild-type CREB5, L434P CREB5, and CREB3. Several canonical AR co-factors (AR, FOXA1, HOXB13) interact with both CREB5 and CREB5 L434P and are shown.

To identify functional interactions specific to wild type CREB5, we performed RIME in LNCaP cells overexpressing V5-tagged CREB5, CREB5 L434P, or luciferase (**Supplemental Table 4**). At approximately the same average peptide counts of CREB5 (8) and L434 CREB5 (7.5), we found a striking correlation between the interaction profiles of CREB5 and CREB5 L434P (R=0.834). This observation indicated that this CREB5 mutant binds the same proteins as the L434P mutant. In parallel, we failed to find a correlation between CREB3 and wild type CREB5 (R=-0.0581) (**Figure 3C**). The limited changes in the L434P CREB5 RIME profile highlighted a subset of differential protein interactions associated with the ART resistant phenotype. When examining the interactions of wild type and L434P CREB5, we found smaller differences between AR, FOXA1 and HOXB13 (**Figure 3D**). Outside of AR co-factors, we detected unique peptide signals at comparable levels with AR, FOXA1 and HOXB13, including NFIC, TBX3 (**Figure 3D**). These exhibited preferential interaction to wild type CREB5 relative to L434P or CREB3. NFIC and TBX3 were also interacted with FOXA1 in our other RIME experiments (**Supplemental Table 3**). These observations provided candidate interactions of CREB5, specifically NFIC and TBX3, that may be essential in ART resistance.

### TBX3 and NFIC are critical for prostate cancer cells and ART resistance

To examine the relative contribution of TBX3 and NFIC toward cell viability, we analyzed genome-scale RNAi screens performed in 8 prostate cancer cell lines as a part of Project Achilles 2.20.1 (Cowley et al., 2014; Shalem et al., 2014; Tsherniak et al., 2017). Upon computing the average dependency scores in DEMETER, we found that NFIC and TBX3 ranked among the strongest dependencies in these prostate cancer cell lines while exhibiting limited overall dependency in the other 495 cell lines (**Figure 4A**). Their relative dependencies were comparable to the strongest gene dependencies, such as FOXA1 and HOXB13, found in previous studies (Pomerantz et al., 2015). While we in parallel examined the CRISPR-Cas9 screens (DepMap, 21Q2), the current prostate cancer cell lines analyzed were limited. To further examine if TBX3 and NFIC were critical in models that were ART resistance, we performed CRISPR-Cas9 mediated gene depletion experiments in a cell line model that spontaneously acquired enzalutamide resistance (Kregel et al., 2016). In this cell line, we expressed 3 distinct sgRNAs that depleted FOXA1 or two sgRNAs that ablated TBX3 or NFIC, we found that the sgRNAs against FOXA1, TBX3 or NFIC all reduced protein levels and decreased viability (**Figure 4B, 4C**). We next examined if TBX3 or NFIC interacted with the key CREB5 co-factor FOXA1 in ChIP-seq experiments performed as part of the Encode project (Consortium, 2012). Based on motif enrichment analyses of the TBX3 or NFIC binding sites, we observed statistically significant enrichment of FOXA1 binding motifs identified in experiments on breast or prostate cancer cell lines (**Figure 4D**). Like CREB5, NFIC also bound B-Zip motifs.

**Figure 4.**
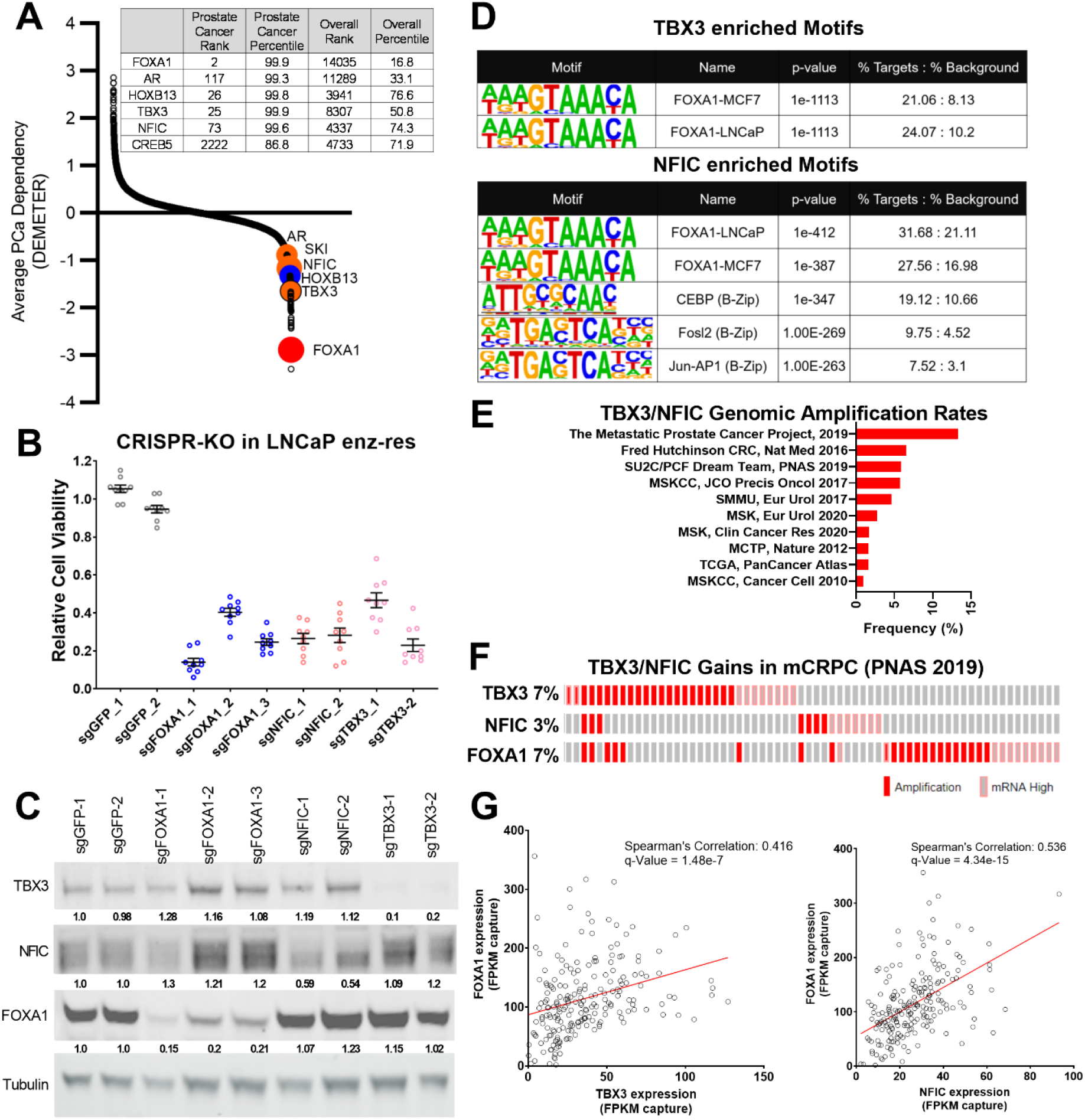
TBX3, NFIC are key regulators in prostate cancer cells including those that are enzalutamide resistant. (A) Analysis of genome-scale RNAi screening data ranking the average dependency of 16869 genes (x-axis) in 8 prostate cancer cell lines and 495 additional cancer cell lines (Project Achilles 2.20.1). Average DEMETER score (y-axis) indicates the dependency correlations of FOXA1 and CREB5-interacting proteins. A negative DEMTER score indicates gene dependency in these specific PC cell lines. Average DEMETER dependency scores for selected genes. (B) CRISPR-Cas9 was utilized to deplete experimental (NFIC, TBX3), negative (GFP) or positive controls (FOXA1) genes in LNCaP cells that spontaneously developed resistance to enzalutamide. The overall cell numbers are depicted post perturbation. (C) A representative immunoblots depicts depletion of proteins from (B) in proliferation experiments. Tubulin was used as a loading control. The relative depletion is quantified based on the average of all experiments after normalizing to tubulin. (D) ChIP-seq data from NFIC and TBX3 was analyzed to predict interaction with CREB5 or FOXA1 motifs. Enriched motifs, the targeted cell lines, and significance levels are depicted. (E) The genomic amplification rates of TBX3 and NFIC are examined in various prostate cancer studies. (F) In one mCRPC study, the rates of TBX3, NFIC and FOXA1 gains are depicted. (G) The expression of TBX and NFIC are compared in one mCRPC study in which the regression line, Spearman’s correlation coefficients and q-values are depicted.

Given their role of regulating FOXA1 functions, we further examined the landscape of TBX3 and NFIC dysregulation in prostate cancer studies using whole exome sequencing and whole transcriptome sequencing data from cBioPortal (Cerami et al., 2012; Gao et al., 2013) based on the 209 mCRPC samples collected by Stand Up 2 Cancer/Prostate Cancer Foundation (SU2C/PCF) (Abida et al., 2019). We found that TBX3 and NFIC genomic amplifications together represented up to 13.3% of prostate cancers, in which notably higher amplification rates were observed in mCRPC studies as opposed to those sampling primary prostate cancer (**Figure 4E**). Upon examining expression and amplification data mCRPC samples (Abida et al., 2019), we found that TBX3, NFIC and FOXA1 were amplified or overexpressed in 7%, 3% and 7% of mCRPC samples, respectively (**Figure 4F**). Of the 209 samples with expression data, we found a robust positive correlation between FOXA1 with TBX3 (Spearman’s correlation: 0.416, q-Value: 1.48e-7) or NFIC (Spearman’s correlation: 0.536, q-Value: 4.34e-15) (**Figure 4G**). Together, we found TBX3 and NFIC were nuclear proteins associated with CREB5, FOXA1 and functionally impacted prostate cancer cell viability and ART resistance even absent of CREB5. In addition, they interacted specifically at FOXA1 motifs in cell lines, are amplified or overexpressed in prostate cancer patients and their expression is associated with that of FOXA1 in mCRPC.

### CREB5 regulates ART resistant pathways in cell lines and mCRPC patients

To determine signaling functions of CREB5, we determined transcriptome expression patterns that were statistically associated with CREB5 expression in both cell line and mCRPC patients (**Figure 5A**). Of the proteins that commonly interacted with CREB5, we utilized Enrichr (Chen et al., 2013; Kuleshov et al., 2016) to perform pathway enrichment analysis on the 335 proteins that bound both CREB5 and FOXA1 in our RIME profiling analysis. Based on the top 10 statistically significant signatures, we found that these 335 interactions associated with AR, cell cycle, as well as Notch, Wnt and SMAD/TGFβ pathways (**Figure 5B**). In clinical mCRPC, we analyzed RNA-sequencing datasets from the SU2C/PCF mCRPC cohort (n = 209) (Abida et al., 2019). We computed the Spearman’s correlation coefficient for *CREB5* expression against each transcriptional target gene in the AR, MYC, and cell cycle signatures. In mCRPC, CREB5 expression was indeed positively correlated with the expression of similar signaling programs we found *in vitro* including epithelial to mesenchymal transition, TGFβ, Wnt, and Notch (**Figure 5C**) (Alumkal et al., 2020; He et al., 2021). Together, the CREB5-associated nuclear protein interactions in cells and transcripts in mCRPC provide mechanistic insights in which ART resistant pathways can be activated by this transcription factor.

**Figure 5.**
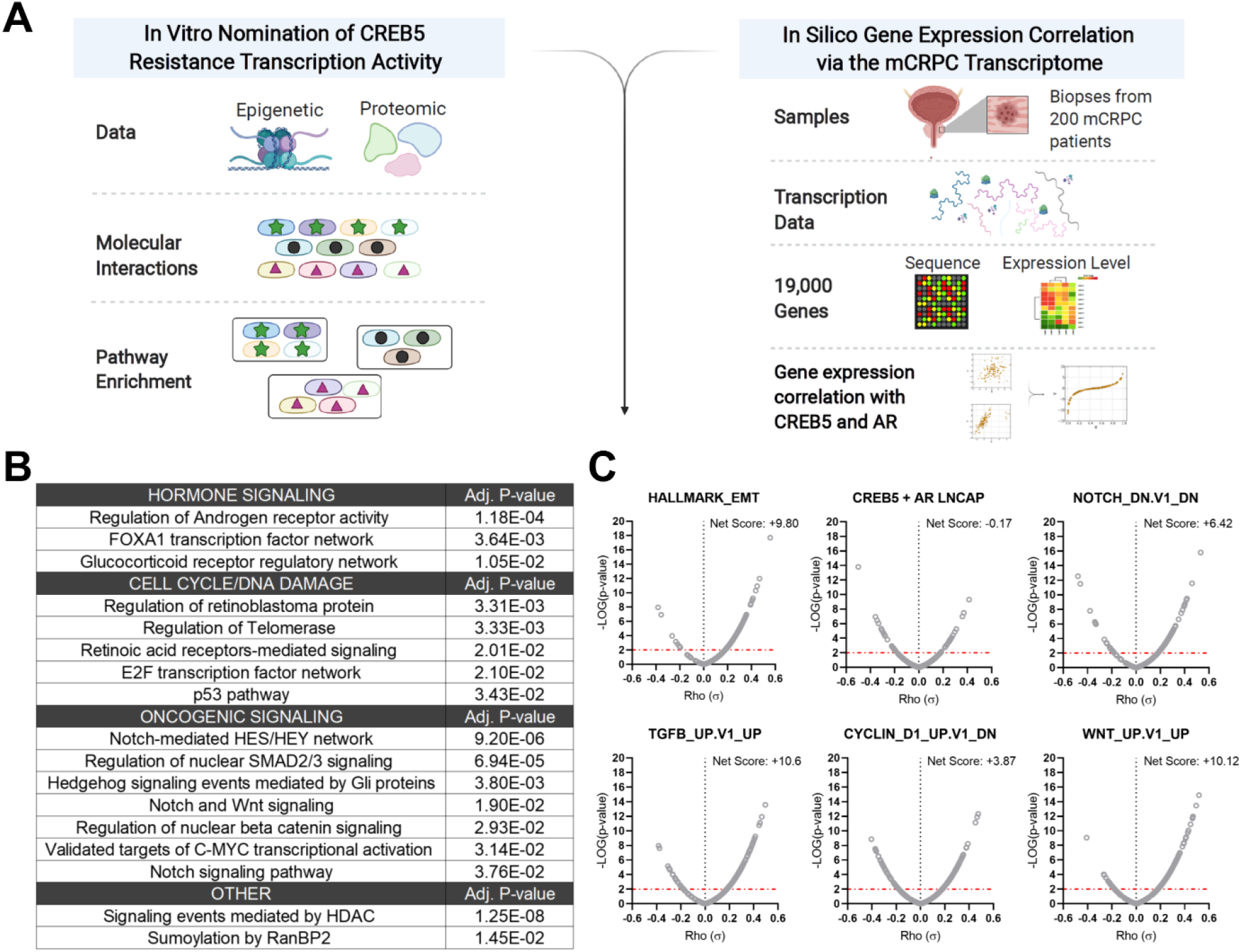
Integrative analysis of CREB5 activity. (A) A workflow of the informatics analysis of CREB5 using *in vitro* and mCRPC data. (B) Spectrum of shared CREB5 and FOXA1 protein interactions identified by RIME are analyzed. The enriched pathways and statistical significance are presented for specific pathways. (C) Based on RNA-seq from clinical mCRPC, Spearman correlation coefficients compare *CREB5* expression with AR, MYC and cell cycle signaling programs. Correlation coefficient values (Rho, σ, x-axis) for *CREB5* against each gene, as represented by a single dot, and the statistical significance (negative log of p-value, y-axis) are displayed. P-value is marked (red dotted line).

## Discussion

We and others have demonstrated transcription regulators, such as FOXA1, promote prostate cancer progression and resistance to ART and ADT. Our work identified other molecular events associated with transition towards resistance to 2^nd^ generation ART and ADT. We did so by identifying reprogramming events associated with overexpressed CREB5, which exhibited shifts in binding at transcription regulatory elements as well as interaction with co-factors when cells were challenged with enzalutamide. In promoting proliferation in ART, CREB5 exhibited a strong degree of interaction with known AR transcription machinery including FOXA1, HOXB13 and GRHL2 as well as novel prostate cancer regulators TBX3 and NFIC. The robust convergence of CREB5-FOXA1 function was observed at through binding of transcription regulatory elements and interactions among nuclear proteins. The factors TBX3 and NFIC interacted with CREB5 but required an intact B/L-Zip domain. TBX3 and NFIC, two nuclear factors that are amplified or overexpressed in mCRPC, were vulnerabilities in other prostate cancer models including ones that were ART resistant, and bound to FOXA1 transcription regulatory elements. Informatics modeling of the CREB5 activity through protein interactions and mCRPC transcription patterns indicated CREB5 is associated with pathways found in patients resistant to enzalutamide (Alumkal et al., 2020; He et al., 2021). Altogether, our study indicates that the dynamic binding properties of CREB5 mediates assembly of essential factors to AR and FOXA1 to promote resistant transcripts (**Figure 6**).

**Figure 6.**
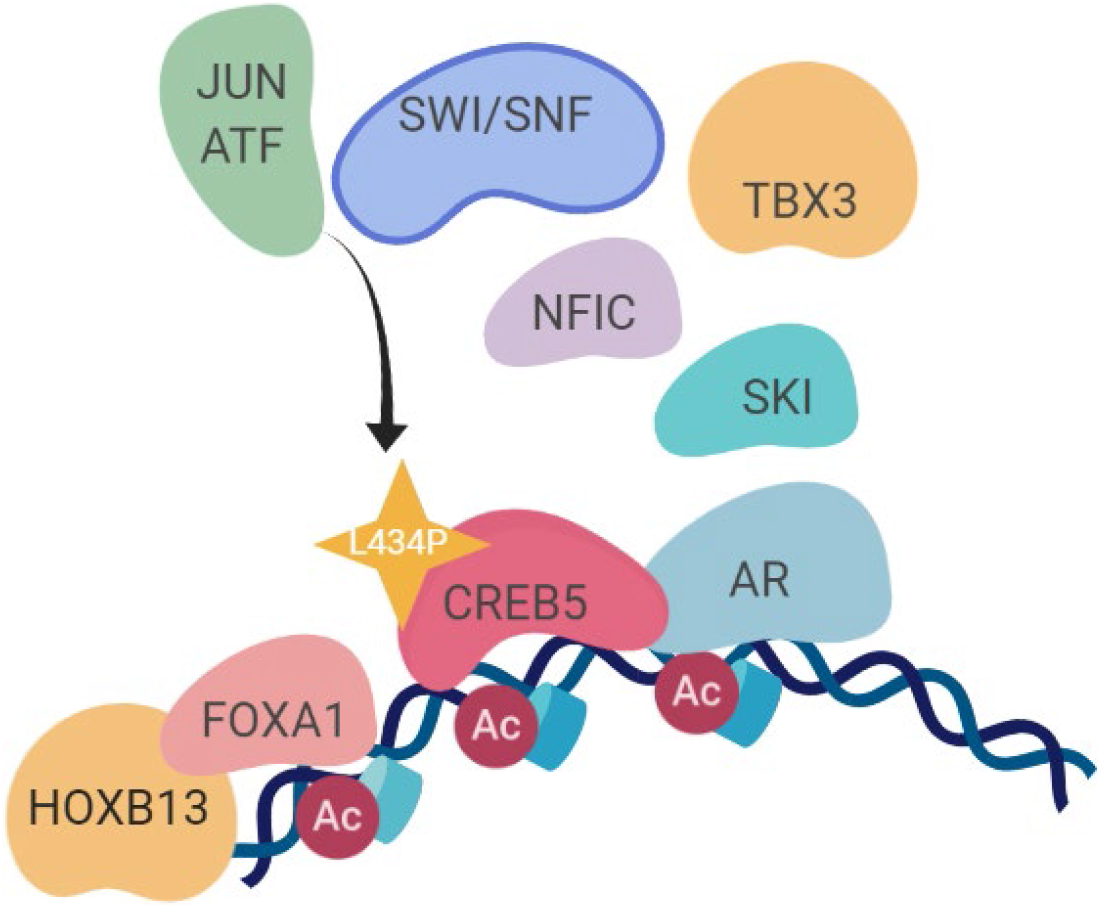
A molecular model of the CREB5 complex and transcription promoting ART resistance.

FOXA1 functions as an oncogenic pioneering and transcription factor in cancers, including prostate and breast (Gerhardt et al., 2012; Nakshatri and Badve, 2009; Shah and Brown, 2019). FOXA1 is a critical dependency in prostate cancer cell line models (Pomerantz et al., 2015), including those that transdifferentiate into AR-agnostic, neuroendocrine like features (Baca et al., 2021). Studies have broadly examined the binding patterns of FOXA1 with transcription regulatory elements in normal prostate tissue, primary prostate cancer, mCRPC, and neuroendocrine prostate cancers (Baca et al., 2021; Pomerantz et al., 2015; Pomerantz et al., 2020), demonstrating transcriptional programs associated with prostate in development, tumor progression and drug resistance. The current study found FOXA1 co-regulators through the CREB5-mediated ART resistance phenotype. Independent of CREB5 and FOXA1, TBX3 expression has been formerly demonstrated to be required for viability of breast cancer cells (Amir et al., 2016; Krstic et al., 2016). NFI factors, NFIA/B/C/X, have previously been shown to interact with FOXA1 to promote the transcription of AR target genes (Grabowska et al., 2014). Future studies that examine FOXA1 interactions in parallel through ChIP-seq and RIME may further elucidate its context specific functions. Particularly, it would be of interest to characterize the universal subset of FOXA1 interactions in prostate cancer tumorigenesis as well as molecular changes associated with mutated forms of FOXA1 (Adams et al., 2019; Parolia et al., 2019).

Studies have identified increases in CREB5 as a marker for metastasis in ovarian, breast and colorectal cancers (Bhardwaj et al., 2017; He et al., 2017; Molnar et al., 2018; Qi and Ding, 2014). Functionally, Bardwaj et al. have identified CREB5 transcripts as a repressed target of miRNA-29c, a tumor suppressive miRNA lost in the triple-negative subtype of breast cancer (Bhardwaj et al., 2017). Forced overexpression of CREB5 promoted cell cycle and colony formation in this study. Altogether, these studies implicate pro-tumor CREB5 functions in cancers. While other CREB family members (Welti et al., 2021) have been associated with therapy resistance in advanced prostate cancer, the differential RIME interaction profiles displayed by these two family member proteins, CREB5 and CREB3, exhibited dichotomous behavior with respect to binding of nuclear proteins in cancer cells. This suggests that the oncogenic roles of CREB5 and other CREB family members may be distinct and mediated through their structurally different N-terminal domains. As another key observation, we also find that experimental conditions including cell culture and drug treatment dramatically influenced CREB5 molecular interactions.

Our findings also support that transcription regulators may act as effective therapeutic targets in mCRPC. As examples, TBX3 and NFCI have been previously detected in large-scale proteomic approaches that interrogated prostate cancer tissue (Sinha et al., 2019). The current study demonstrates they have key regulatory roles in prostate cancer cell viability and ART resistance. Antagonizing nuclear or transcription factors has been efficacious in recent examples, as inhibitors against EP300, an AR interacting protein, were efficacious in prostate cancer models (Jin et al., 2017; Lasko et al., 2017; Welti et al., 2021). In regulating a pro-tumorigenic role in mCRPC, CREB5 require additional factors (**supplemental table 2, 3, 4**) including JUN/ATF and SWI/SNF complex. While we have previously discussed that CREB5 functions differ from other CREB or ATF family members (Hwang et al., 2019), the context of how CREB5 interacts with CREB or ATF factors, such as potential hetero-dimerization at open regions of chromatin, still requires further resolution through biochemical approaches. Of bound SWI/SNF family members, we have previously discussed SMARCB1 as recurrent mutation in mCRPC (Armenia et al., 2018), and its loss as a biomarker in malignant pediatric tumors (Hong et al., 2019; Howard et al., 2019). How CREB5 interacts with other chromatin remodeling complexes in ART resistance is an additional research direction that could improve our understanding of transcription processes that could act as targets in therapy resistant mCRPC.

In summary, our observations implicate CREB5 as a driver of mCRPC. At the molecular level, our findings depict a complex model of therapy resistance that occurs in the nucleus of tumor cells that permits the activation of oncogenic signaling pathways. Furthering the understanding of these underlying changes may inform of additional research avenues and precision strategies for advanced cancer patients that depend on CREB5.

## Acknowledgements

This work was supported in part by the Weizmann Institute of Science – National Postdoctoral Award Program for Advancing Women in Science (to R.A), Targets of Cancer Training program grant T32 CA009138 (to M.L.), Ray of Light Foundation (to H.E.B.), NIH/NCI (K00 CA212221) (To J.P.R.), American Cancer Society-AstraZeneca (PF-16-142-01-TBE) (to J.H), Young Investigator Award from the American Society of Clinical Oncologists (ASCO) and by the PhRMA Foundation and Kure It Cancer Research Foundation (To S.C.B), U01 CA233100 (E.M.V.), Mark Foundation Emerging Leader Award (E.M.V.), U.S. National Institutes of Health/National Cancer Institute: U01 CA176058 (to W.C.H.).

We acknowledge Joshua Pan from Dana Farber Institute and Broad Institute of MIT and Harvard for designing approaches for co-dependency analysis. Graphical figures were created with BioRender.com.

